# Androgen Signaling in Type 2 Innate Lymphoid Cells Drives Sex Differences in *Helicobacter*-Induced Gastric Inflammation and Atrophy

**DOI:** 10.1101/2025.03.14.643321

**Authors:** Benjamin C. Duncan, Maeve T. Morris, Jordan L. Pascoe, Stuti Khadka, Lei Wang, Gangqing Hu, Jonathan T. Busada

**Author notes:** Corresponding authors: Jonathan T. Busada, WVU School of Medicine, Microbiology, Immunology and Cell Biology, 64 Medical Center Drive, P.O. Box 9177, Morgantown, WV 26506.

## Abstract

**Background & Aims:** Gastric cancer is the fifth most common cancer worldwide. Men are disproportionately affected by gastric cancer, which ranks as the fourth most common cancer in men compared to eighth in women worldwide. Chronic inflammation driven by *Helicobacter pylori* infection remains the leading gastric cancer risk factor. Emerging evidence suggests that sex hormones modulate immune responses, contributing to sex differences in infection outcomes and cancer susceptibility. This study investigates how androgens influence the gastric inflammatory response to *Helicobacter* infection and contribute to sex disparities in disease progression.

**Methods:** Male and female C57BL/6 mice were colonized with *Helicobacter felis* to investigate sex differences in gastric inflammation. Androgen levels were manipulated by bilateral castration in males and dihydrotestosterone (DHT) treatment in females. Single-cell RNA sequencing was used to identify androgen-responsive leukocyte populations and to establish cell communication networks between leukocyte clusters. The functional roles of these cells were further defined using ILC2- and T cell-deficient mouse models.

**Results:** Infected female mice developed significantly more severe gastric inflammation, atrophy, and metaplasia infection compared to males. Androgen depletion by castration increased gastric inflammation and accelerated preneoplastic lesion development, while these pathological features were reduced by DHT treatment. Androgen-responsive type 2 innate lymphoid cells (ILC2s) were key initiators of gastric inflammation and ILC2 depletion abolished the sex differences in *H. felis* pathogenesis.

**Conclusions:** This study reveals that androgens suppress *Helicobacter*-induced gastric inflammation by modulating ILC2 activation. We found that androgens are protective, as androgen depletion exacerbated gastric inflammation and accelerated preneoplastic lesion development. These findings provide mechanistic insight into the age-related increase in male gastric cancer incidence, coinciding with declining androgen levels. Our results suggest that circulating androgen concentrations may serve as a prognostic biomarker for gastric cancer risk in men.

**Graphical Abstract:** 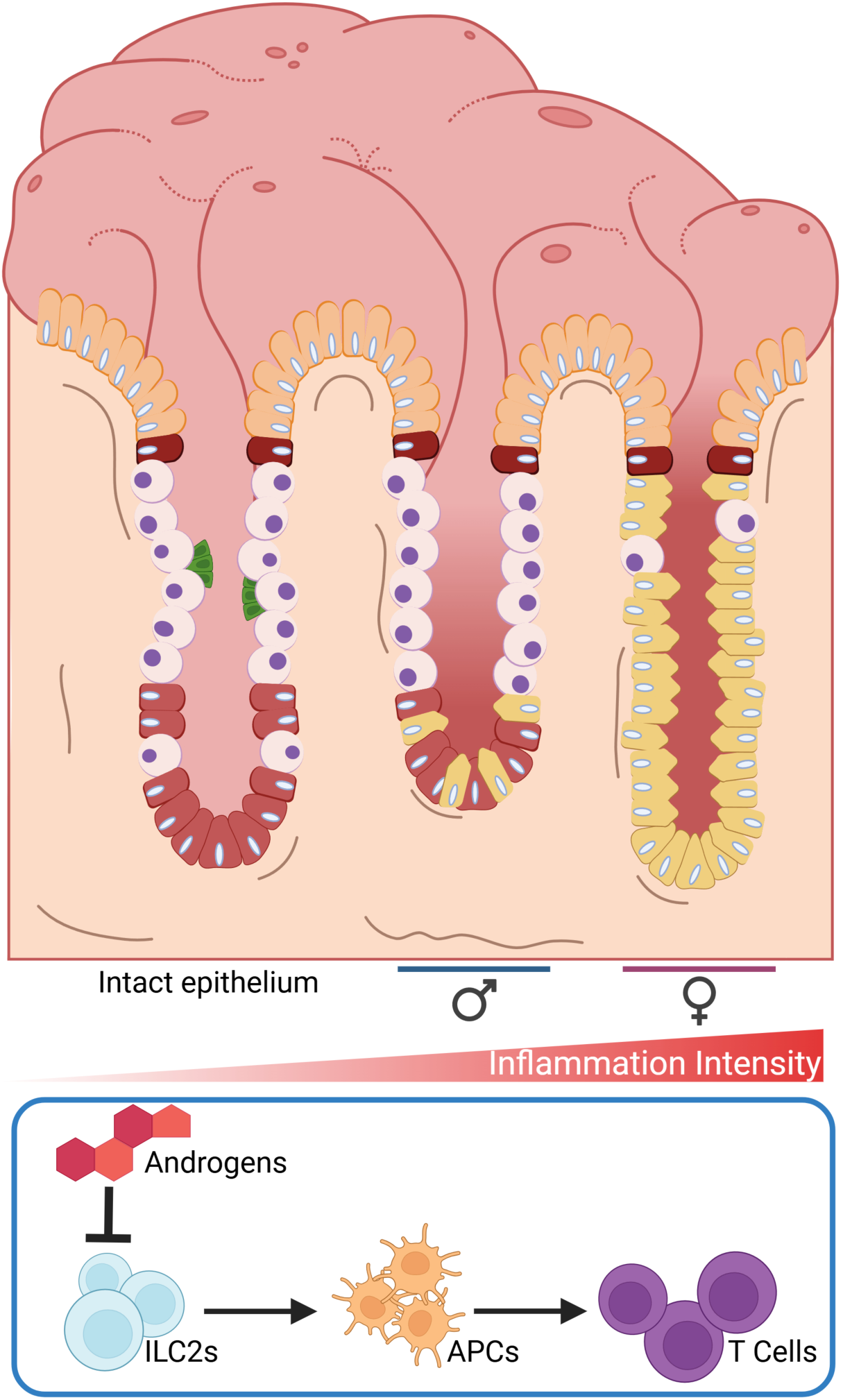

## Introduction

Sex steroid hormones act as important rheostats for controlling sex differences in immune function. While their effects on the immune system are pleiotropic, estrogens tend to be pro-inflammatory by enhancing effector T cell responses and stimulating cytokine production ^1^. In contrast, androgens are predominantly anti-inflammatory, suppressing innate and adaptive immune responses and inhibiting the transcription of genes encoding pro-inflammatory cytokines ^2^. Extensive experimental and clinical evidence supports that females mount stronger and more effective immune responses to infections and vaccines than males ^1-3^. Consequently, females are generally better protected from infectious diseases but are more susceptible to hypersensitivity and autoimmune disorders. In contrast, men tend to exhibit increased mortality from infections and higher rates of cancer.

Gastric cancer is the fifth most common cancer worldwide ^4^. Infection by the Gram-negative bacteria *Helicobacter pylori* remains the leading gastric cancer risk factor and is associated with up to 90% of non-cardia gastric cancer cases ^5^. Chronic inflammation is a critical driver of *H. pylori* pathogenesis and strong Th1/Th17-polarized immune responses drive gastric atrophy and preneoplastic lesion development, while tolerogenic immune responses are protective ^6-8^. Despite the link between intense immune responses and gastric carcinogenesis, gastric cancer is a profoundly male-biased disease, with more than 65% of global gastric cancer cases diagnosed in men ^4^. While lifestyle factors such as smoking and alcohol consumption contribute to male susceptibility, the sex disparity persists even after adjusting for these risks ^9, 10^, suggesting that endogenous biological factors – such as sex hormones, influence gastric cancer risk.

Type 2 innate lymphoid cells (ILC2s) are tissue-resident leukocytes in the gastric mucosa. Upon epithelial damage, ILC2s stimulate gastric macrophage recruitment and IL13 production, promoting pyloric metaplasia (PM) maturation ^11-14^. While ILC2s were initially described for their role in producing type 2 cytokines such as IL5 and IL13 in response to damage-associated molecular patters (DAMPS), they are now recognized as highly plastic, capable of producing mixed Th1/Th2-associated cytokines during bacterial infections ^15, 16^. We have previously demonstrated that androgens suppress gastric ILC2 function by regulating the transcription of proinflammatory cytokines, such as *Il13* and *Csf2 ^11^*. Similar findings have been made in the pulmonary system, where androgen suppression of ILC2s protects from hypersensitivity reactions ^17, 18^. Recently, ILC2s were also shown to control sex differences in the skin in response to *Staph epidermidis* infection, directing antigen presenting cell (APC) and T cell recruitment and activation ^19^. In the stomach, *H. pylori* infection increases the number of ILC2s, where they contribute to humoral immune responses ^20^. However, their role in *H. pylori-* associated atrophic gastritis and subsequent metaplasia remains unclear.

Gastric cancer is the most common pathogen-associated malignancy, and chronic inflammation is required to foster its development. Paradoxically, androgens tend to suppress inflammation in males, gastric cancer rates are higher in men than in women. Interestingly, this sex difference in gastric cancer rates does not emerge until middle age. Data from the National Cancer Institute’s Surveillance, Epidemiology, and End Results (SEER) program show similar gastric cancer rates in young men and women, with a dramatic increase in men after the age of 50 ^21^. This increase coincides with age-related declines in androgen levels, known as andropause (or male menopause) ^22^. Thus, androgens may protect young males from *H. pylori-*driven inflammation, and androgen decline in middle age may accelerate carcinogenesis. However, the mechanisms by which host sex influences *H. pylori*-driven immune responses and subsequent disease progression remain poorly understood. In this study, we define how host sex shapes *Helicobacter-*driven gastric inflammation and subsequent preneoplastic lesion development. Our findings reveal that males are significantly protected from *Helicobacter*-associated pathologies. We identified a novel ILC2 – APC – T cell axis that controls sex differences in *Helicobacter-* induced gastric inflammation. Moreover, we provide evidence that androgens suppress gastric inflammation, as androgen depletion by castration significantly increased inflammation and accelerated preneoplastic lesion development in males. Collectively, our results demonstrate a critical role for ILC2s in shaping sex differences in the gastric response to *Helicobacter* infection and suggest that androgens levels may serve as a prognostic marker for assessing *H. pylori* pathogenesis.

## Material and Methods

### Animal Care and Treatment

All mouse studies were performed with approval by the West Virginia University Animal Care and Use Committee. C57BL/6J, TCRβδKO (strain number 002122), ROSA-DTA (strain number 009669) and Red5 (strain number 030926) mice were purchased from the Jackson Laboratory. Mice were administered standard chow and water *ab libitum* and maintained in a temperature- and humidity-controlled room with standard 12-hour light/dark cycles. The TCRβδKO strain was maintained in sterile housing for immunocompromised mice. The ROSA-DTA and Red5-Cre were crossed to constitutively deplete IL5-expressing cells. Mice with the genotype ROSA-DTA; *Il5* Cre+/- were considered ILC2 deficient (called ILC2 DTx hereafter) while mice with the genotype ROSA-DTA; *Il5* Cre -/- were used as WT littermate controls. 8-week-old mice were used for all experiments. Mock mice received 500μL sterile brucella broth, and infected mice were inoculated with 500 μL of brucella broth containing 10^9^ CFU of the bacteria two times 24 hours apart as previously described ^23^. Castration and drug pellet implantation were performed as previously described ^11^.

### Bacteria Preparation

*H. felis* (ATCC 49179) was grown on tryptic soy agar plates (BD Biosciences) with 5% defibrinated sheep blood (Hemostat Labs) and 10 µg/mL vancomycin (Alfa Aesar) under microaerophilic conditions (5% O_2_ and 10% CO_2_) at 37°C for 2 days. *H. felis* was then harvested and transferred to Brucella broth (Research Products International) containing 5% fetal bovine serum (R&D Systems) and 10 µg/mL vancomycin and grown overnight at 37°C under microaerophilic conditions with agitation. Bacteria were centrifuged and resuspended in fresh Brucella broth without antibiotics before spectrophotometry and mouse infection.

### Tissue Preparation

Stomachs were opened along the greater curvature and washed in phosphate buffered saline to remove the gastric contents. Stomachs used for histology were pinned to cork boards and fixed overnight in 4% paraformaldehyde at 4°C. For histology, strips were taken from the greater curvature of the gastric corpus. Stomach strips were then transferred into 70% ethanol and submitted to the West Virginia University histology core for routine processing, embedding, sectioning, and H&E staining. Images were obtained using an Olympus VS-120 brightfield slide scanning microscope. Stomachs used for immunostaining were obtained from the lessor curvature and cryopreserved in 30% sucrose and then embedded in optimal cutting temperature media.

### Immunohistochemistry

Immunostaining was performed using standard methods. Briefly, 5-μm stomach cryosections were incubated with anti H+/K+ ATPase antibodies (clone 1H9, MBL Life Science), MIST1 (clone D7N4B; Cell Signaling Technologies), CD45 (clone 104; Biolegend), CD68 (clone FA-11; Biolegend) or CD44v9 (Cosmo Bio) for 1 hour at room temperature or overnight at 4°C. Sections were incubated in secondary antibodies for 1 hour at room temperature. Fluorescence-conjugated *Griffonia simplicifolia* lectin (GSII; ThermoFisher) was added with secondary antibodies where indicated. Sections were mounted with Vectastain mounting media containing 4’,6-diamidino-2-phenylindole (Vector Laboratories). Images were obtained using a Zeiss 710 confocal laser-scanning microscope (Carl-Zeiss) and running Zen Black (Carl-Zeiss) imaging software. Parietal cells and chief cells were quantitated as previously described ^24^ using confocal micrographs captured with a ×20 microscope objective. Cells were counted using the ImageJ count tool (National Institutes of Health). Cells that stained positive with anti-H^+^-K^+^ antibodies were identified as parietal cells and cells that stained positive with anti-MIST1 antibodies and were GSII negative were identified as mature chief cells. Cell counts were normalized to image area to determine the cell number per 100 μm^2^. Images that contained gastric glands cut longitudinally were selected for counting.

### RNA Isolation and qRT-PCR

Gastric tissue RNA biopsies were collected from the greater curvature using a 2-mm biopsy punch. RNA was extracted in TRIzol (Thermo Fisher Scientific) and precipitated from the aqueous phase using 1 volume of 100% ethanol. The mixture was transferred to an RNA isolation column (Omega Bio-Tek) and the remaining steps were followed according to the manufacturer’s recommendations. RNA was treated with RNase-free DNase I (Omega Bio-Tek) as part of the isolation procedure. Reverse transcription followed by qPCR was performed in the same reaction using the Universal Probes One-Step PCR kit (Bio-Rad Laboratories) and the TaqMan primers/probe mixtures (all from Thermo Fisher Scientific): *Ppib* (Mm00478295_m1), *Aqp5* (Mm00437578_m1), *Gkn3* (Mm01183934_m1).

### Flow Cytometry

Corpus tissue from euthanized mice was washed in Hanks Balanced Salt Solution without Ca^2+^ or Mg^2+^ containing 5 mM HEPES, 5 mM EDTA, and 5% FBS at 37°C plus 1μl/ml dithiothreitol. Tissue was then washed in Hanks Balanced Salt Solution with Ca^2+^ or Mg^2+^ and then digested in 1mg/ml collagenase (Worthington Biochemical). After digestion, the tissue fragments were pushed through a 100 μM strainer and centrifuged. Pelleted cells were resuspended and rinsed through a 40 μM strainer. Intact cells were enriched using an Optiprep (Serumwerk) density gradient. Fc receptors were blocked with TruStain (Biolegend) and then stained with antibodies. The following antibodies were used: CD45.2 (clone 104), CD3e (clone 145-2C11), B220 (clone RA3-6B2), CD4 (clone GK1.5), CD8a (clone 53-6.7), CD11b (clone M1/70), MHCII (clone M5/114.15.2) Ly6g (clone 1A8), F4/80 (clone BM8), SiglecF (clone E50-2440), Lineage cocktail (Biolegend), CD127 (clone A7R34), Ly-6A/E Sca-1 (clone D7), ICOS (clone 15F9). Actinomycin D (clone A1310, Invitrogen) was used to label dead cells. Super Bright Complete staining buffer (Invitrogen) was used in antibody cocktails containing multiple brilliant violet fluorophores. Cells were analyzed on a Cytek Aurora spectral flow cytometer (Cytek Biosciences). Flow cytometry analysis was performed using Cytobank analysis software (Beckman Coulter).

### RNAseq

Mock-infected and *Helicobacter felis*-infected male and female mice were euthanized two months post-inoculation. Stomachs were harvested, and RNA was isolated as previously described. RNA libraries were prepared using a NEB Ultra II Directional RNA Library Prep Kit with poly-A selection (Admera Health). Sequencing was performed on a paired-end platform, generating 150 bp reads with a depth of 40 million reads per sample. Following sequencing, the West Virginia University Bioinformatics Core processed the raw data to generate a list of differentially expressed genes (DEGs) and a reads per kilobase per million (RPKM) file. The DEG list included log2 fold change, adjusted *p*-values, and other statistical metrics. These data served as the input for subsequent Gene Set Enrichment Analysis (GSEA).

### GSEA

Gene Set Enrichment Analysis (GSEA) was performed using the clusterProfiler and REACTOMEPA packages in R to identify significantly enriched pathways associated with immune-related processes. All truly differentially expressed genes were ranked by log2 fold changes (logFC) derived from differential gene expression analysis. Both mouse-specific KEGG and REACTOME gene sets were utilized for the analysis.

For KEGG pathway analysis, the gseKEGG function was applied with the following parameters: a minimum gene set size of 10, a maximum size of 500, and a significance threshold of adjusted p-value < 0.05. REACTOME pathway analysis was conducted using the gsePathway function in the REACTOMEPA package. Parameters included a minimum gene set size of 10, a maximum size of 500, and a significance threshold of adjusted p-value < 0.05. The KEGG database and REACTOME database-specific annotations were mapped to mouse-specific Entrez IDs.

Visualization of enriched pathways was performed using ridge plots generated with the enrichplot package. The plots illustrate the distribution of ranked gene-level statistics for genes within each pathway, highlighting their contributions to pathway enrichment. A color gradient was used to denote the Normalized Enrichment Score (NES). Specific pathways of interest were selected based on their biological relevance.

### Single-cell RNAseq

Previously published Single-cell RNAseq dataset were used for this study ^11^. H5 files were downloaded from the NCBI Gene Expression Omnibus, GSE147177 and imported into R (version 4.3.3). The files contain scRNA seq data from pooled (n=4) CD45+ leukocytes isolated from the gastric corpus of normal males, normal females, and adrenalectomized+castrated males. Gene expression matrices were converted into a Seurat object using the Seurat R package (version 5.0.3). Gene expression measurements were normalized using the Seurat NormalizeData function and standardized using ScaleData. Next, genes that expressed high variability across cells were identified using Seurat’s FindVariableFeatures function. The Seurat RunPCA function and visualization of the components via elbow plot were performed and the first 20 components were selected for further analyses. The RunUMAP, dimensions = 20, min.distance = .5 and n.neighbors= 20, function was then applied and the FindNeighbors function was used using the first 20 dimensions to construct a Shared Nearest Neighbor Graph. FindClusters with “resolution = .5” was performed to cluster cells into respective clusters. All marker genes with a minimum .4 pct expression value were identified using Seurat’s FindAllMarkers function and exported to an excel table. Additionally, UMAP plots were converted to a loupe file using 10XGenomics Loupe Browser using the create_loupe_fromSeurat function. Annotation and color visualization was performed in Loupe Browser (version 8.0.0). Marker genes were identified by the average expression value in the cluster of interest and had a minimum pct value of .8 or higher while the minimum pct was below .4. This insured high expression of the gene profile within the cluster while limiting the expression in the other clusters. Once clusters were identified, gene matrices were subsetted using the subset.data function to include only the clusters of interest for further analysis.

### Communication Networks

Subsetted Seurat objects containing the gene expression matrices were converted into a cell chat object using CellChat’s (version 2.1.2) createCellChatobject function. CellChat’s identifyOverExpressedGenes function was used to find overexpressed genes in clusters of interest using CellChat’s mouse data bank. Ligand-receptor pairs were identified using CellChat’s identifyOverExpressedInteractions to further analyze cell communication. A communication probability matrix was generated using the computeCommunProb function using the TriMean to calculate probability scores. Once calculated, communication networks were visualized using CellChat’s netVisual_circle function.

### Statistical Analysis

All error bars are ± SD of the mean. The sample size for each experiment is indicated in the figure legends. Experiments were repeated a minimum of 2 times. Statistical analyses were performed using 1-way analysis of variance with the post hoc Tukey t-test when comparing 3 or more groups or by an unpaired t-test when comparing two groups. Statistical analysis was performed by GraphPad Prism 10 software. Statistical significance was set at *P* ≤ 0.05. Specific *P* values are listed in the figure legends.

## Results

### Males mount blunted inflammatory responses to *Helicobacter* infection compared to females

Chronic inflammation is a key driver of *H. pylori* pathophysiology, promoting gastric atrophy and metaplasia development and shaping gastric cancer risk ^25^. Females tend to mount more robust inflammatory responses to infection compared to males ^1^. To investigate sex differences in *Helicobacter-*induced gastric inflammation, we utilized the well-established *Helicobacter felis* model, which rapidly induces gastric inflammation, atrophy, and metaplasia in mice ^23^. Male and female C57BL/6J mice were colonized by *H. felis,* and the stomachs were collected 2 months post-colonization. Evaluation by CD45 immunostaining and flow cytometry indicated that male and female mock-infected mice did not exhibit significant sex differences in gastric-resident immune cells (Figure 1A-B). *H. felis* colonization significantly increased leukocyte infiltration into the gastric corpus in both sexes. However, leukocyte infiltration was significantly blunted in male mice compared to females (Figure 1A-B). Immunophenotyping by flow cytometry (see Supplementary Figure 1 for gating strategy) revealed that macrophage, T cell, and eosinophil infiltration were significantly blunted in *H. felis-*infected males compared to females (Figure 1B and Supplementary Figure 2A), while B cell and neutrophil recruitment were equivalent between both sexes (Supplementary Figure 2A). Within the T cell compartment, males were significantly biased towards a CD4+ T cell response, while females were biased toward increased CD8+ T cell recruitment (Supplementary Figure 2B). To further characterize sex differences in the gastric response to *H. felis,* we performed bulk RNA sequencing on the gastric corpus of infected males and females. *H. felis* colonization resulted in 608 differentially expressed genes in males while inducing 2806 differentially expressed genes in females (Supplementary Figure 3). Next, GSEA was utilized to identify genetic signatures and pathways enriched in males and females. GSEA-enriched pathways from the KEGG and REACTOME databases were visualized by Ridge plot, revealing broader pathway activation and higher enrichment scores of immune-related gene networks in infected females compared to males, particularly in pathways governing adaptive immune responses, antigen presentation, and myeloid cell function (Figure 1C). Together, these data demonstrate that females mount a stronger gastric immune response to *Helicobacter* infection compared to males.

**Figure 1.**
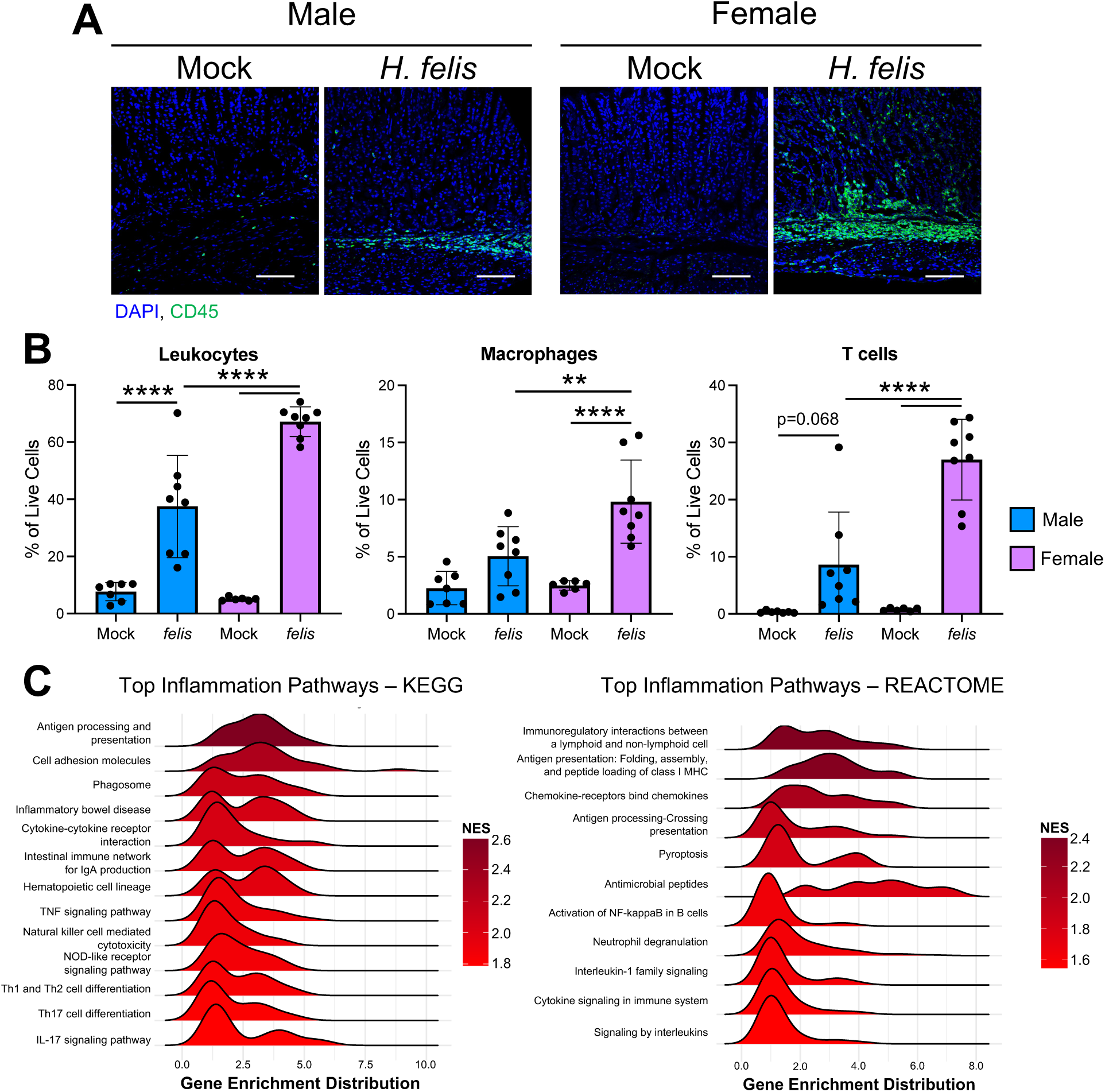
Females mount more intense gastric inflammatory responses to *H. felis* infection. (A) Representative immunofluorescent staining of CD45+ immune cells (green) in the gastric corpus of male and female mice. Nuclei are counterstained with DAPI (blue). Scale bars: 100 µm. (B) Flow cytometry quantification of the indicated leukocyte populations in the gastric corpus of male and female mice. *p < 0.05, **p < 0.01, ***p < 0.001, ****p < 0.0001. n≥6. (C) Ridge plots display of Gene Set Enrichment Analysis (GSEA) pathway enrichment. The plots visualize the distribution of ranked gene-level statistics for genes within each enriched pathway, with the density curve representing the relative distribution of genes in the ranked list. n=3.

### Males are protected from *Helicobacter-*driven gastric atrophy and metaplasia development

Chronic inflammation during *Helicobacter* infection drives gastric atrophy and metaplasia development ^26^. Given our results demonstrating that males mount blunted immune responses to *H. felis* compared to females, we next assessed *Helicobacter-*driven gastric atrophy and metaplasia development. There were no discernible histological differences between mock-infected males and females (Figure 2A). *H. felis-*infected male mice developed modest chief cell and parietal cell atrophy (Figure 2A-C). In contrast, female corpus glands exhibited severe remodeling and mucous neck cell hyperplasia, denoted by *Griffonia Simplicifolia II* lectin staining, as well as severe chief and parietal cell atrophy, with near complete loss of both cell types. (Figure 2A-C). To assess sex differences in PM development, we screened the bulk RNAseq DEG list described above for a panel of well-described PM-associated transcripts (Supplementary Table 1). Among these 14 PM markers, all were significantly expressed in infected females compared to infected males (Figure 2D). To confirm these results, qRT-PCR was performed for the PM-transcripts *Aqp5* and *Gkn3,* both of which were significantly increased in *H. felis-*infected females but were not significantly elevated in infected males (Figure 2E). Finally, immunostaining for the *de novo* pyloric metaplasia marker CD44v9 revealed scattered PM-positive gastric units within the male gastric corpus (Figure 2F), whereas CD44v9 staining was widespread in *H. felis-*infected females. In parallel with increased inflammation, these data demonstrate that *Helicobacter-*infected females develop more severe atrophy and PM compared to infected males.

**Figure 2.**
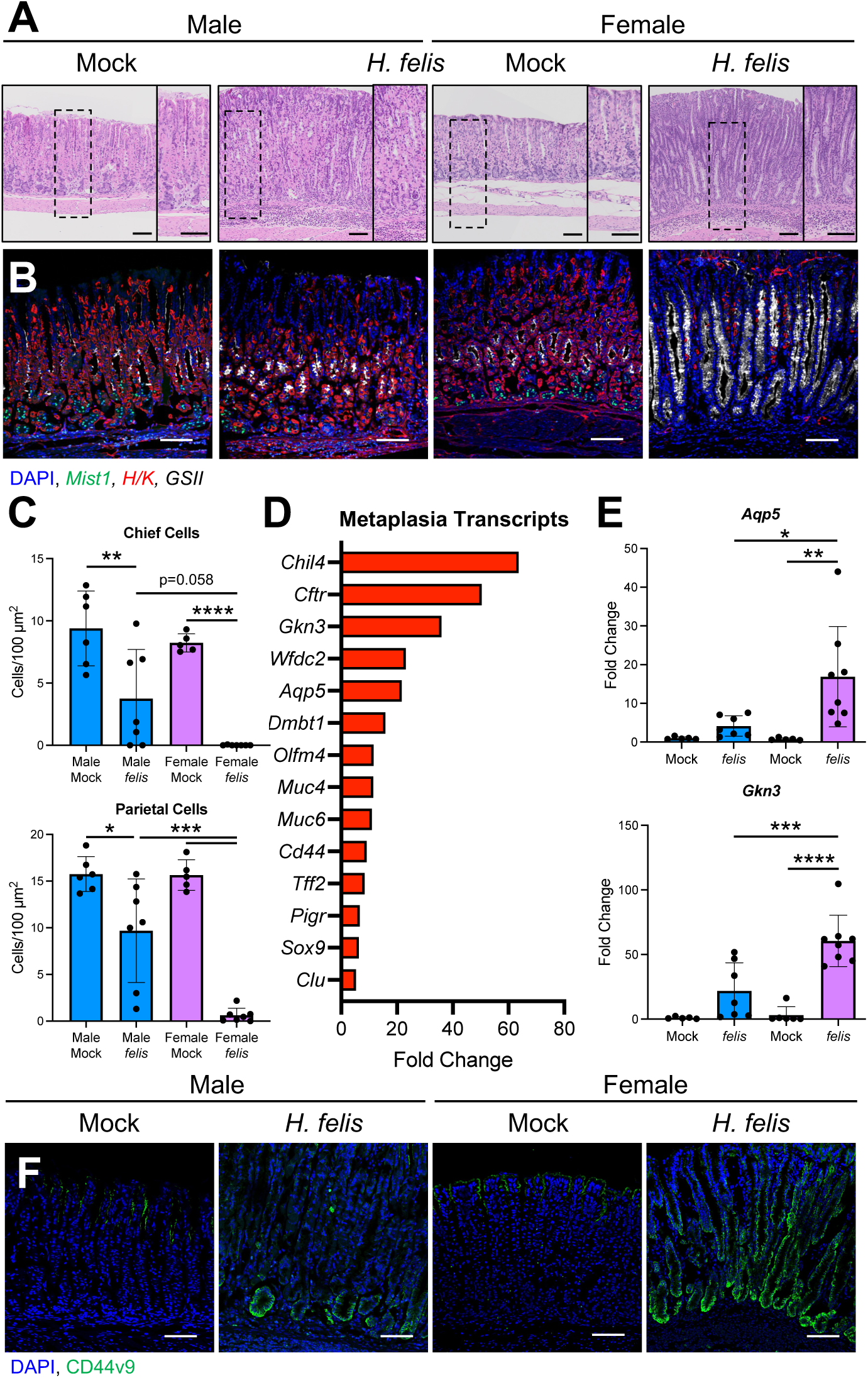
Males are protected from *H. felis-*driven gastric atrophy and metaplasia. (A-B) Representative micrographs of (A) H&E-stained sections or (B) immunostaining of H+/K+ ATPase (red, parietal cells), MIST1 (green, chief cells), GSII lectin (white, mucous neck cells), and DAPI (nuclei, blue) of gastric mucosa from male and female mice 2 months post *H. felis* colonization. (C) Quantification of chief and parietal cell loss between males and females. (D) Fold change of metaplasia-associated genes. The bar plot displays the top-ranked genes associated with metaplasia, ranked by fold change. Gene expression was analyzed and ranked based on differential expression and p-value. (E) qRT-PCR of the metaplasia transcripts *Aqp5* and *Gkn3*. (F) Immunofluorescent staining of the metaplastic marker CD44v9 (green) in the gastric corpus. All scale bars = 100 µm. *p < 0.05, **p < 0.01, ***p < 0.001, ****p < 0.0001. n≥6, except D where n=3.

### Sex differences in *Helicobacter* pathogenesis are controlled by androgens

Sex hormones play important roles in regulating immune function and androgens predominately oppose immune activation ^2^. We have previously shown that androgen signaling protects the stomach from inflammatory damage ^11^. To investigate the physiological mechanisms underlying male protection from *Helicobacter* infection, we castrated adult male mice prior to *H. felis* colonization. Two months post-inoculation, immunostaining and flow cytometry revealed that castrated *H. felis-*infected mice displayed a significant increase in leukocyte infiltration within the gastric corpus compared to intact-infected males (Figure 3A-B). Moreover, castration significantly increased gastric macrophage and T cell recruitment (Figure 3B), which are critical effectors of *Helicobacter-*driven gastric atrophy and PM development ^24, 26^. We next evaluated how castration impacted *H. felis-*driven gastric atrophy and metaplasia development. H&E micrographs demonstrated that castration dramatically accelerated *H. felis* histopathological progression with increased mucous neck cell hyperplasia and gastric atrophy (Figure 3C-D). Quantification of immunostaining revealed that castration significantly increased chief and parietal atrophy (Figure 3D-E). Next, qRT-PCR for *Aqp5* and *Gkn3* demonstrated that these PM markers were significantly increased in castrated mice compared to intact infected male controls (Figure 3F). Similarly, immunostaining demonstrated a dramatic increase in CD44v9+ gastric units (Figure 3G). These data demonstrate that castration enhances *H. felis-*driven gastric inflammation, atrophy, and metaplasia and suggest that male sex hormones contribute to sex differences in *Helicobacter* pathogenesis.

**Figure 3.**
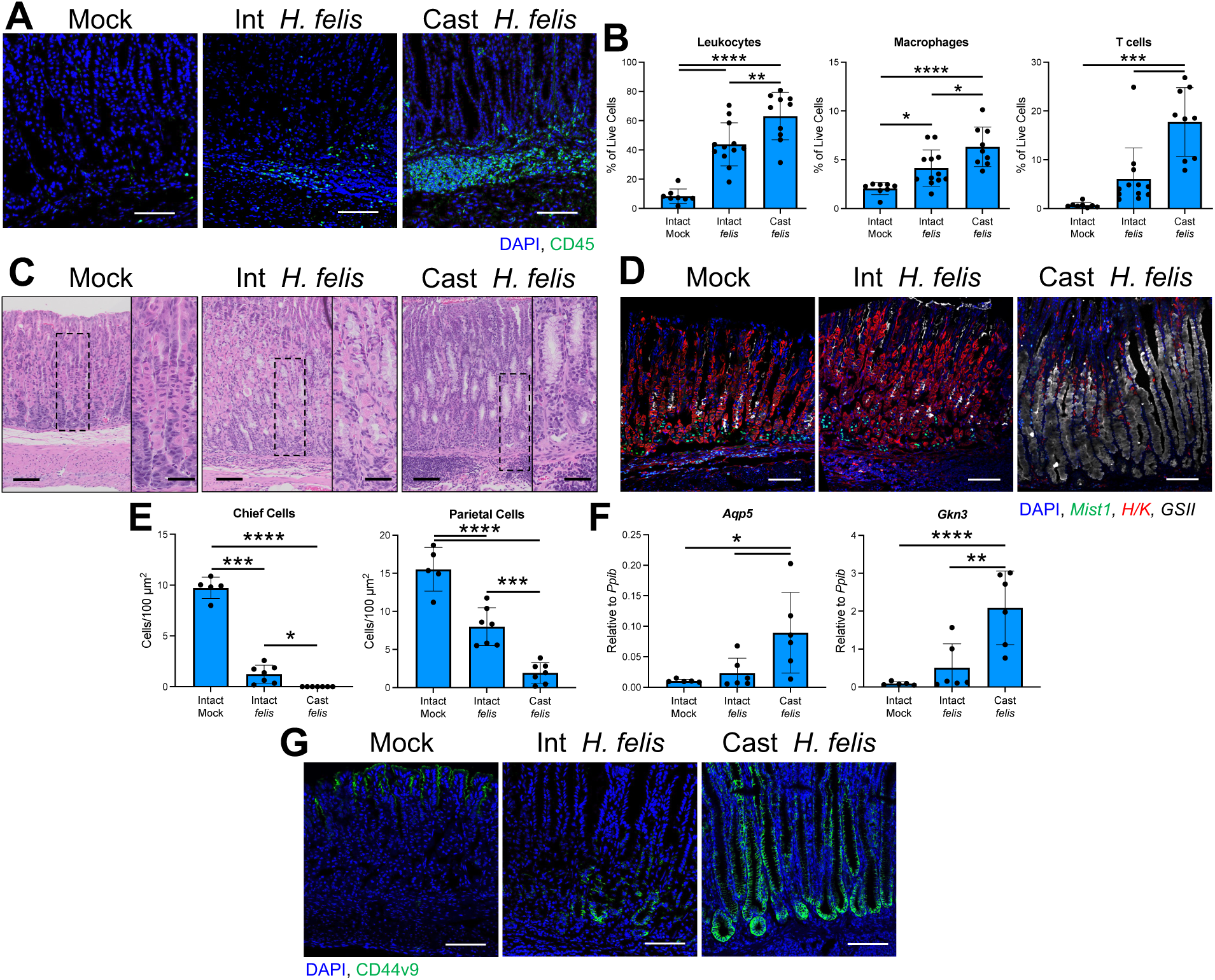
Androgen depletion via castration enhances gastric inflammation and accelerates preneoplastic lesion development. (A) Representative immunofluorescent staining of CD45+ immune cells (green) in the gastric corpus of male and female mice. Nuclei are counterstained with DAPI (blue). (B) Flow cytometry quantification of the indicated leukocyte populations in the gastric corpus of intact male and castrated mice 2 months post *H. felis* colonization. (C-D) Representative micrographs of (C) H&E-stained sections or (D) immunostaining of H+/K+ ATPase (red, parietal cells), MIST1 (green, chief cells), GSII lectin (white, mucous neck cells), and DAPI (nuclei, blue) of gastric mucosa from intact male or castrated male mice. (E) Quantification of chief and parietal cell loss. (F) qRT-PCR of the metaplasia transcripts *Aqp5* and *Gkn3*. (G) Immunofluorescent staining of the metaplastic marker CD44v9 (green) in the gastric corpus. Scale bars = 100 µm and 50 µm in the insets (C). *p < 0.05, **p < 0.01, ***p < 0.001, ****p < 0.0001. n≥5.

We next investigated whether androgens can protect from *Helicobacter-*induced gastric inflammation, atrophy, and metaplasia by surgically implanting a 5⍺ dihydrotestosterone (DHT) time-release pellet into female mice prior to inoculation with *H. felis* (Figure 4A). DHT treatment significantly suppressed CD45+ immune cell infiltration compared to placebo-treated *H. felis* infected controls (Figure 4B-C). Immunophenotyping revealed that DHT treatment significantly suppressed the recruitment of gastric macrophages and T cells (Figure 4C). We next evaluated how DHT treatment impacted *H. felis-*driven gastric atrophy and metaplasia development. H&E micrographs demonstrated gross remodeling of the gastric mucosa in female *H. felis* infected placebo controls, while remarkably, the stomach of DHT-treated females appeared qualitatively normal (Figure 4D). Immunostaining quantification revealed that DHT treatment significantly prevented chief and parietal atrophy compared to placebo-treated controls (Figure 4D-E). Next, qRT-PCR for *Aqp5* and *Gkn3* demonstrated that these PM markers were significantly increased in placebo-treated infected controls, while their expression did not significantly increase in infected DHT-treated mice (Figure 4F). Similarly, CD44v9 immunostaining was widespread in placebo-treated *H. felis-*infected females, while staining was undetected in infected females treated with *H. felis* (Figure 4G). These results demonstrate that androgen suppresses the gastric immune response to *H. felis* and protects from subsequent gastric atrophy and PM development.

**Figure 4.**
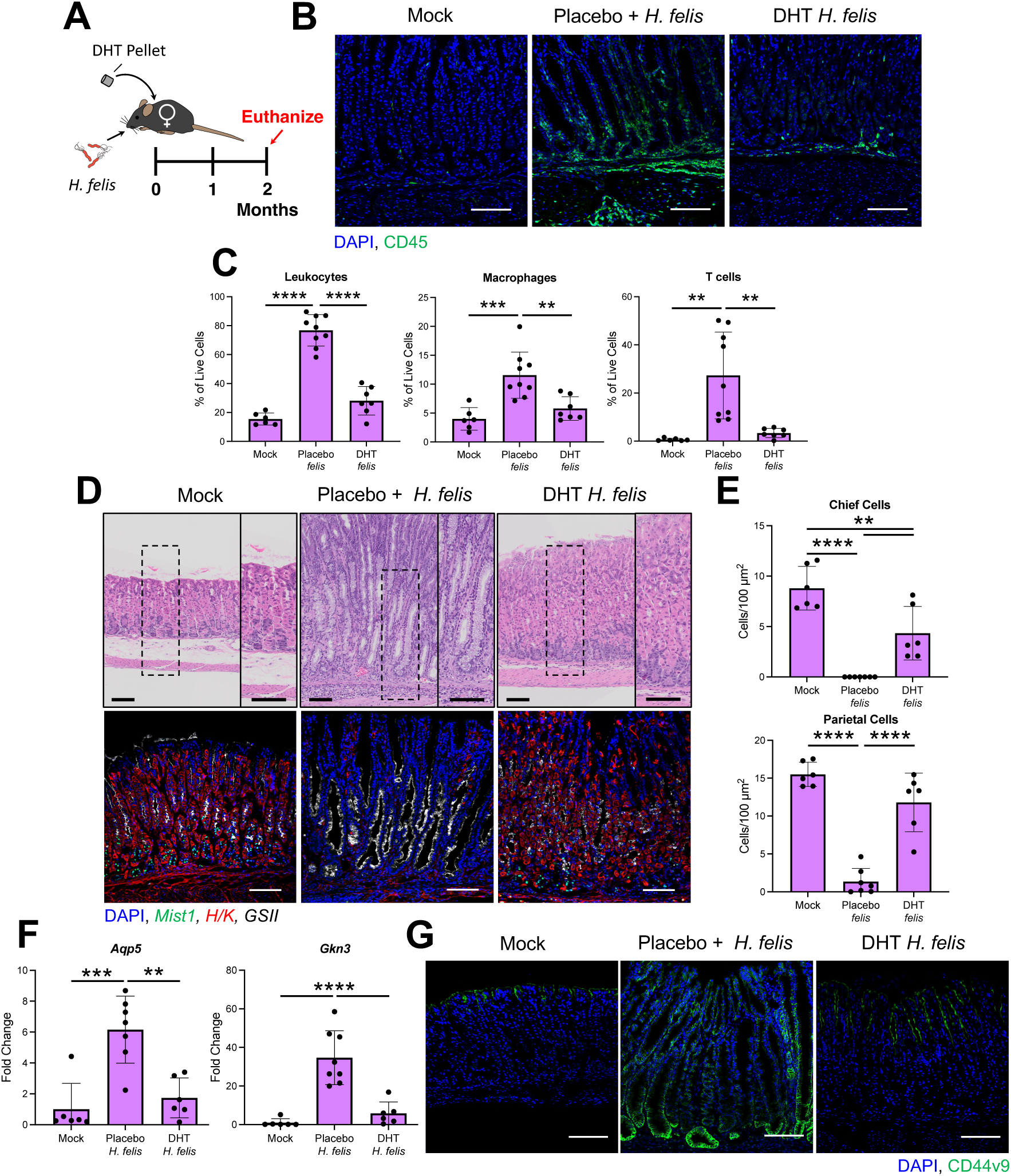
DHT suppresses *H. felis* inflammation, atrophy, and metaplasia development. (A) Schematic representing experimental design. (B) Representative immunofluorescent staining of CD45+ leukocytes (green) in the gastric mucosa of mock-inoculated, *H. felis-*infected placebo-treated, and DHT-treated female mice. Nuclei are counterstained with DAPI (blue). (C) Flow cytometry quantification of the indicated leukocyte populations in the gastric corpus. (D) Representative micrographs of (top) H&E-stained sections or (bottom) immunostaining of H+/K+ ATPase (red, parietal cells), MIST1 (green, chief cells), GSII lectin (white, mucous neck cells), and DAPI (nuclei, blue) from the gastric corpus. (E) Quantification of chief and parietal cell loss. (F) qRT-PCR of the metaplasia transcripts *Aqp5* and *Gkn3*. (G) Immunofluorescent staining of the metaplastic marker CD44v9 (green) in the gastric corpus. All scale bars = 100 µm. **p < 0.01, ***p < 0.001, ****p < 0.0001. n≥6.

### Androgen responsive ILC2s coordinate the gastric inflammatory response to *Helicobacter* infection

Our results demonstrated that androgens suppress the gastric immune response to *H. felis* infection, reducing the recruitment of macrophages and T cells. Androgens primarily signal through the androgen receptor (AR). Therefore, we analyzed our previously described scRNA seq of gastric leukocytes sorted from the gastric corpus of male, female, and castrated male mice to identify *Ar*+ leukocyte populations ^11^. The cells were dimensionally reduced into nine clusters using the Uniform Manifold Approximation and Projection (UMAP) algorithm (Figure 5A). Each cluster was annotated by unique marker signatures, which were depicted in a dot plot (Figure 5B). The relative expression of *Ar* transcript density per cell cluster was depicted by violin plot. Three cell *Ar*+ cell clusters—ILC2s, T cells, and NK cells—were identified, with ILC2s displaying the most ubiquitous expression (Figure 5C). Interestingly, the *Ar* was not expressed within the macrophage or dendritic cell clusters, in agreement with our previous findings ^11^, despite macrophage recruitment being enhanced following castration and suppressed by DHT treatment (Figures 3B and Figure 4B, respectively). These results suggest that an androgen-responsive cell type directs macrophage recruitment. Therefore, we mapped communication networks between gastric ILC2, the APCs macrophages and dendritic cells, and T cells leukocytes using CellChat. Cell communication scores between ILC2s and APC were dramatically lower in males compared to females, while scores between APC and T cells were similar between both sexes (Figure 5D). Castration dramatically increased the score between ILC2s and APCs, while the scores between APC and T cells remained largely unchanged. These results suggest that ILC2s regulate an APC – T cell axis within the stomach and that androgen signaling suppresses communication between ILC2s and APCs.

**Figure 5.**
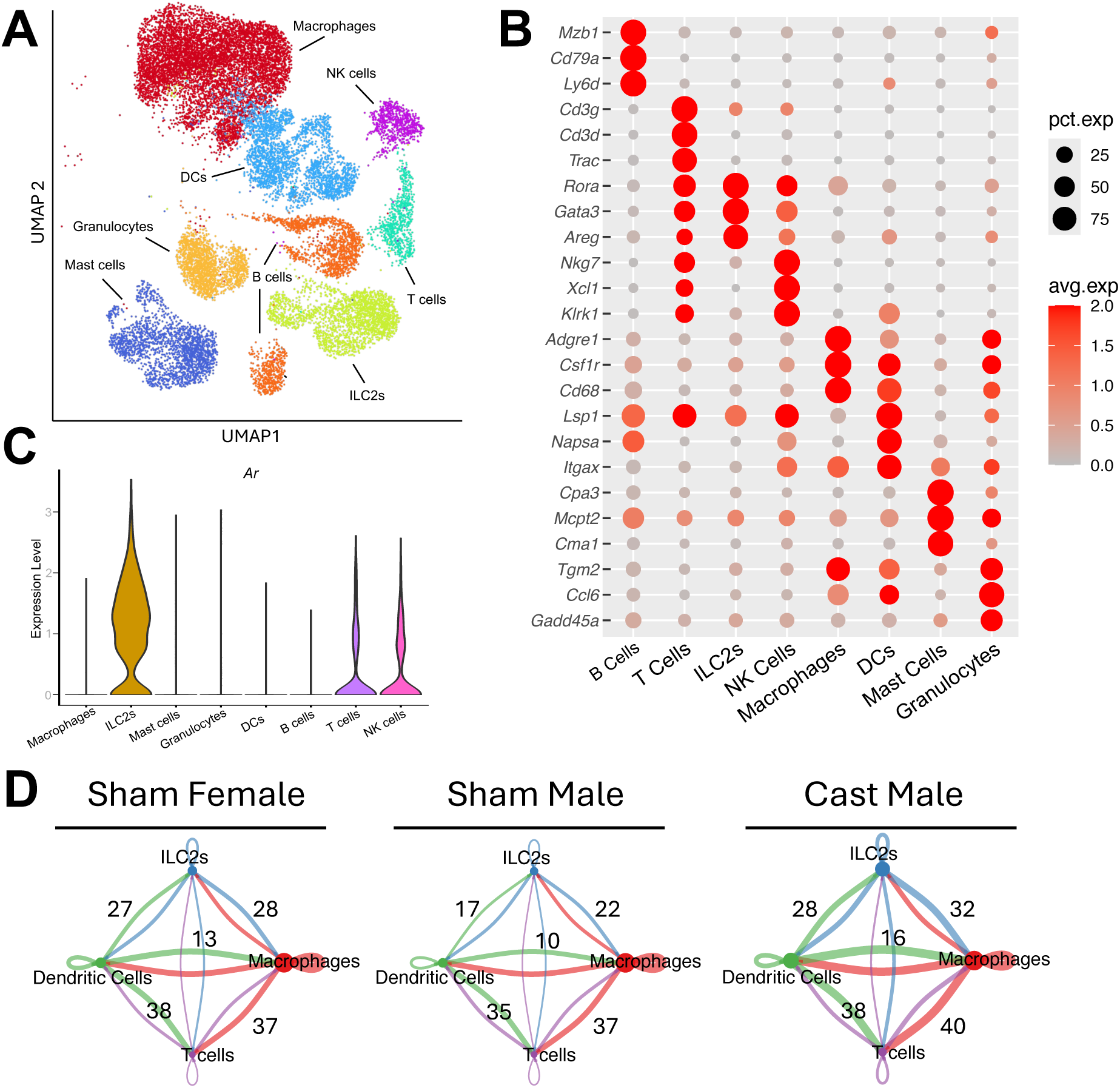
Single-cell RNA communication networks predict and androgen responsive ILC2-APC-T cell axis within the gastric immune milieu. (A) UMAP plot of immune cell clusters within the gastric corpus of pooled sham female, sham male, and castrated male mice (B) Dot plot showing the expression of key marker genes used to identify immune cell populations. Dot size represents the percentage of cells expressing the gene within each cluster, and dot color indicates the average expression level. (C) Violin plot of androgen receptor (*Ar*) expression across immune cell populations (D) Intercellular communication networks predicted using CellChat. Lines represent ligand-receptor interactions between immune cell populations, with line thickness indicating the strength of interaction. Cell were pooled from 4 individual mice for each group.

### ILC2s initiate an APC—T cell an axis during *Helicobacter* infection

Next, we sought to define the leukocyte communication axis that regulates the gastric inflammatory response to *Helicobacter* infection. Based on our scRNA seq results, we hypothesized that ILC2s regulate the recruitment of macrophages and T cells during *H. felis* infection (Figure 6A). We employed a genetic approach to deplete either ILC2s or T cells to test this hypothesis. Our scRNA seq demonstrated that *Il5* is highly restricted to gastric ILC2s (Figure 6B). Therefore, we selected the Red5 *Il5* Cre driver to target ILC2s ^27^. While *Helicobacter* infection is predominantly Th1/Th17 polarized, IL5 is also expressed by Th2 polarized T cells ^28-30^. Therefore, Red5 Cre mice were colonized with *H. felis* and gastric leukocytes were collected 2 months post-infection. Flow cytometry analysis revealed that tdTomato was highly expressed within gastric ILC2s and was not detectable within gastric T cells (Figure 6 C-D). Next, Red5-Cre mice were crossed with a ROSA-DTA Cre-inducible diphtheria toxin to deplete ILC2s (ILC2-DTx). In this mouse model, Cre removes a stop codon, activating diphtheria toxin expression, which induces cell death. Flow cytometry was used to measure gastric ILC2s in ILC2s-DTx mice and WT littermate controls, revealing a >4.5-fold reduction in gastric ILC2s (Figure 6E and Supplementary Figure 4).

**Figure 6.**
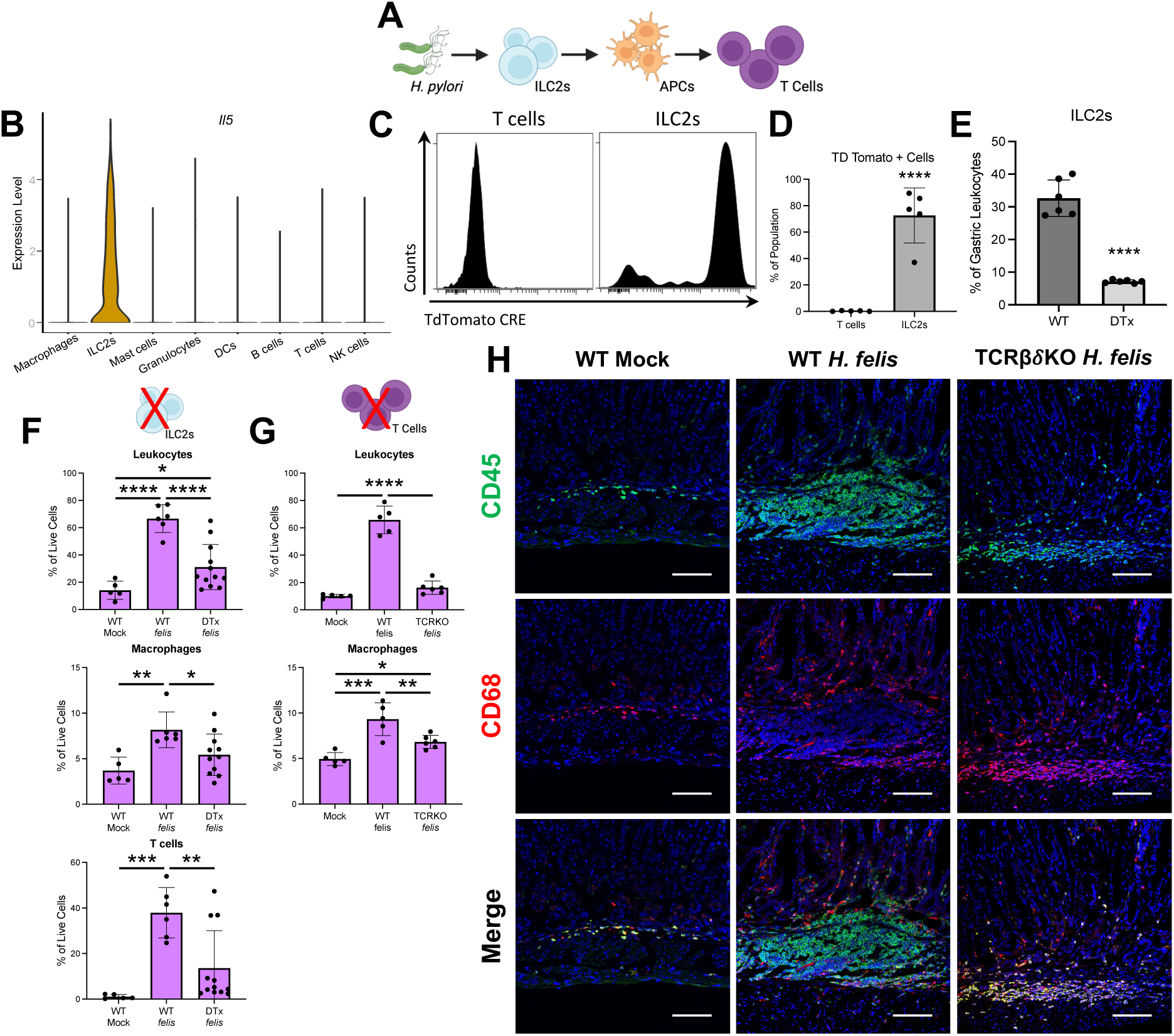
ILC2s control the recruitment of macrophages and T cells during *H. felis* infection. (A) Schematic depicting putative ILC2 – APC – T cell axis. (B) Violin plot of *Il5* expression across immune cell populations. (C-D) Flow cytometry depicting IL5-tdTomato fluorescence in T cell and ILC2s isolated from Red5-Cre mice 2 months after colonization with *H. felis*. (E) Flow cytometry quantification of gastric ILC2s in non-infected WT and ILC2 DTx mice. (F-G) Flow cytometry quantification of the indicated leukocyte populations in (F) TCRBD double KO mice or (G) ILC2-DTx mice 2 months after *H. felis* colonization. (H) Representative immunostaining staining of CD45+ leukocytes (green) and the macrophage marker CD68 (red) in the gastric mucosa of TCRB double KO mice. Scale bar: 100 µm. (*p < 0.05, **p < 0.01, ***p < 0.001, ****p < 0.0001) n≥5.

Next, to define how ILC2 depletion impacted *H. felis-*induced gastric inflammation, ILC2-DTx stomachs were assessed by flow cytometry 2 months following *H. felis* colonization. Female ILC2-DTx mice exhibited a significant reduction of total gastric leukocyte infiltration compared to WT littermate controls (Figure 6F). Moreover, gastric macrophage and T cell infiltration was significantly reduced, suggesting that ILC2s are required to recruit these cell types following *H. felis* colonization. Next, to evaluate the impact of gastric T cell depletion, T cell receptor (TCR) beta/delta double KO mice – which lack all mature T cells – were colonized with *H. felis* and collected 2 months post-infection. Flow cytometry revealed that total gastric leukocyte infiltration was significantly blunted in TCRβδ KO mice (Figure 6G). However, gastric macrophage infiltration remained significantly elevated. Immunofluorescence imaging indicated while macrophages were only a subset of gastric leukocytes in infected WT controls, nearly all gastric leukocytes were CD68+ macrophages in TCRβδ mice (Figure 6G-H). These results suggest that gastric ILC2s are critical for initiating gastric macrophage and T cell recruitment following *Helicobacter* colonization.

### ILC2 depletion protects from *Helicobacter-*induced gastric atrophy and metaplasia

ILC2s produce large quantities of Th2-associated cytokines, such as IL13, that are well-known to drive pyloric metaplasia development ^12, 14^. We have previously reported that androgens suppress cytokine production in ILC2s ^11^. Therefore, we next assessed how ILC2 depletion impacted *H. felis-*induced gastric atrophy and metaplasia development. Stomachs from DTx mice were collected 2 weeks (acute) and 2 months (chronic) post-*H. felis* inoculation. Acute *H. felis* infection induced significant chief and parietal cell atrophy in WT female mice (Figure 7A-C). In contrast, *H felis-*infected ILC2-DTx mice were grossly histologically normal and did not exhibit significant changes in the number of chief or parietal cells (Figure 7A-C). Chronic *H. felis* infection induced significant chief and parietal cell atrophy in both WT and ILC2-DTx mice, but parietal cell atrophy was significantly blunted in the ILC2 DTx mice compared to WT infected controls (Figure 7A-C). ILC2 depletion had no overt impact on *H. felis* pathogenesis in male mice, which remained histologically indistinguishable from infected WT males at both 2 weeks and 2 months post-infection (Supplementary Figure 5), suggesting that, in males, ILC2s are already functionally suppressed. Next, PM development was assessed by qRT-PCR for *Aqp5* and *Gkn3* and by immunostaining for CD44v9. WT infected controls exhibited significant induction of these PM markers at both 2 weeks and 2 months post-infection (Figure 7D-E). In contrast, increased expression of PM markers was completely ablated in LC2-DTx mice following acute *H. felis* infection remained significantly reduced following chronic infection (Figure 7D-E). Finally, immunostaining for CD44v9 demonstrated abundant PM+ gastric units in WT mice (Figure 7F). In contrast, no CD44v9 staining was detected in ILC2-DTx mice 2 weeks post-infection and scattered PM+ units were present 2 months post-infection. These data demonstrate that ILC2 depletion protects from *H. felis* induced gastric atrophy and metaplasia development.

**Figure 7.**
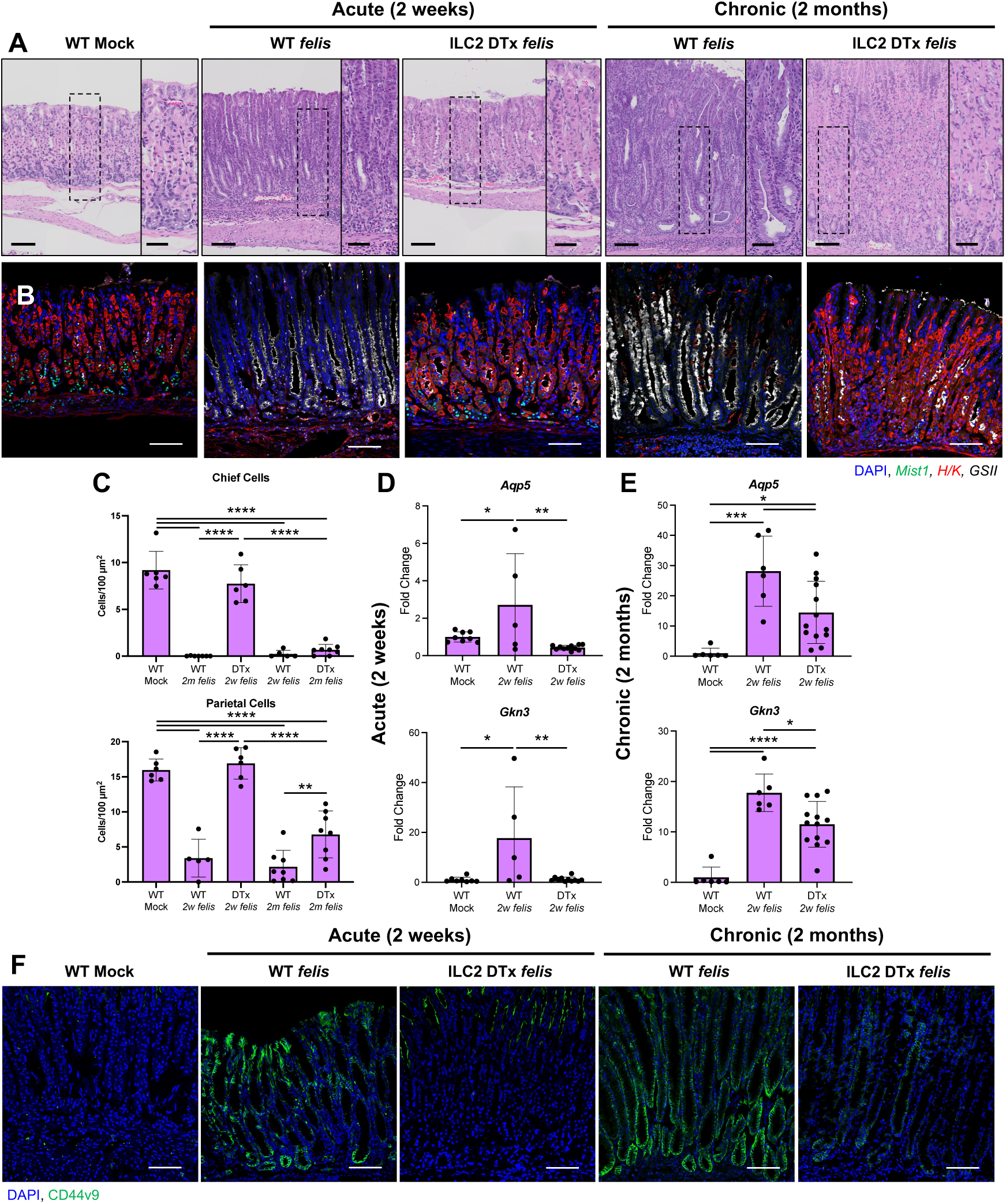
ILC2 depletion protects against *H. felis-*induced gastric atrophy and metaplasia. (A) Representative micrographs of H&E staining of the gastric corpus of acute (2 week) and chronic (2 month) infected wild type (WT) and ILC2-deficient (DTx) mice. (B) Representative immunofluorescent staining of corpus tissue probed with H+/K+ ATPase (red, parietal cells), MIST1 (green, chief cells), and GSII lectin (white, mucous neck cells) in the gastric mucosa. Nuclei are counterstained with DAPI (blue). Scale bar: 100 µm. (C) Quantitation of chief and parietal cell loss between WT and ILC-DTx females at 2 weeks and 2 months per 20x field. (D-E) qRT-PCR analysis of the metaplastic markers *Aqp5* and *Gkn3* in ILC2-DTx mice (D) two weeks and (E) two months post *H. felis-*colonization. (F) Immunofluorescent staining of the metaplastic marker CD44v9 (green). All scale bar = 100 µm. (*p < 0.05, **p < 0.01, ***p < 0.001, ****p < 0.0001). n≥5.

## Discussion

*H. pylori* remains one of the most prevalent chronic bacterial infections worldwide, affecting approximately 44% of the world population ^31^. Colonization by *H. pylori* triggers chronic mucosal inflammation within the gastric mucosa, driving gastric atrophy, metaplasia, and, if untreated, may progress to gastric adenocarcinoma. Chronic inflammation plays a central role in *H. pylori* pathogenesis, and studies using experimental models clearly demonstrate that chronic inflammation is both necessary and sufficient for driving gastric cancer development ^26, 32-34^. However, the causative relationship between severe gastric inflammation and gastric cancer appears to conflict with well-documented sex differences in immune function.

Sex-based differences in immunity are key determinants of infectious disease progression and neoplasia development. Mounting evidence from clinical and experimental studies support that males mount weaker immune response to infectious agents, vaccines, and neoplasia’s compared to females ^3, 35^. While this robust immune activity in females confers greater protection from infections and reduces cancer incidence, it also increases their susceptibility to autoimmune diseases ^36^. Sex hormones significantly contribute to these sex differences in immune function. Here, we demonstrate significant sex differences in the gastric immune response to *H. felis* infection. Our findings agree with a previous report that *H. felis-*infected females developed more widespread gastritis, epithelial proliferation and apoptosis compared to males ^37^. Our results highlight the need of defining how biological sex and sex hormones shape *H. pylori* pathogenesis. While some clinical studies have reported a male bias in *H. pylori-* associated gastritis and peptic ulcer disease, they do not account for confounding variables such as environmental exposures and lifestyle factors that may influence these sex disparities ^38, 39^. Behaviors such as smoking and alcohol consumption and occupational exposures impact *H. pylori* pathogenesis and are more prevalent in men ^40, 41^. A recent study that controlled for sex differences in smoking and alcohol consumption reported significantly higher levels of monocyte infiltration and a trend toward more severe atrophic gastritis in women ^42^. These findings underscore the importance of controlling for lifestyle factors in future studies exploring sex differences in *H. pylori* infection and disease progression.

Most non-reproductive cancers are male-predominant ^43^, a trend that is particularly exacerbated in gastric cancer, with >65% of global gastric cancers diagnosed in men ^4^. While *H. pylori* infection is the leading gastric cancer risk factor, a variety of lifestyle and behavioral factors likely contribute to the male predominance of this disease. However, the sex disparity persists even after adjusting for these variables, suggesting that endogenous biological factors, such as sex hormones, also play a role ^9, 10^. Androgens, have been implicated in the male predominance of a host of tumor types ^35^, but their role in gastric cancer remains unclear. In the stomach, androgens have been shown to exacerbate epithelial damage in ulcer models and impair wound healing ^44, 45^. Paradoxically, our findings reveal that androgens suppress *Helicobacter-*driven gastric inflammation, protecting against gastric atrophy and metaplasia development. Findings from epidemiological studies on androgens and gastric cancer are also mixed. Some studies suggest that anti-androgen therapies reduce gastric cancer incidence ^46^, while others associate higher androgen levels with reduced risk ^47-49^. Interestingly, while total gastric cancer incidence is higher in men, this sex difference does not emerge until middle age. Data from the SEER program indicate that gastric rates are similar in young men and women but dramatically increase in men after the age of 50 ^21^. his increase coincides with the age-related decline in androgen levels—known as andropause ^22^, raising the possibility that declining androgens levels accelerate gastric tumorigenesis. Our results support this notion, demonstrating that androgen treatment protects from *H. felis-*driven gastric inflammation, while androgen depletion via castration significantly increases inflammation and accelerates preneoplastic lesion development. These findings suggest that androgen levels influence gastric cancer risk and may serve as a prognostic biomarker in *H. pylori*-infected men.

Our findings reveal critical roles for ILC2s in coordinating the gastric inflammatory response to *Helicobacter* infection. While previous studies have shown that gastric ILC2s promote PM development and maturation in pathogen-free models of inflammation ^12, 13, 50^, our results demonstrate a broader role for ILC2 in detecting bacterial colonization and initiating immune responses. We found that ILC2 depletion effectively abolished acute gastric inflammation following *H. felis* colonization, significantly reducing macrophage and T cell recruitment.

Although the exact mechanisms by which ILC2s regulate leukocyte recruitment remain unclear, recent studies show that ILC2-derived GM-CSF regulates APC and T cell activation in the skin ^19^. Similar mechanisms may operate in the stomach, as we previously showed that androgens regulate *Csf2* (which encodes GM-CSF) expression in ILC2s, and that gastric macrophages and dendritic cells highly express the GM-CSF receptors *Csf2ra* and *Csf2rb* ^11^.

PM is considered to be a preneoplastic lesion that arises in response to gastric epithelial damage ^51^. Previous studies have shown that ILC2-derived IL13 is required for PM development and maturation ^12, 14^. We have previously reported that DHT suppresses *Il13* expression within ILC2s ^11^. In this study, we found that while ILC2 depletion significantly delayed PM development; however, PM-positive gastric units began to appear at 2 months post-infection, suggesting that alternative sources of IL-13 may contribute to PM formation in the absence of ILC2s. Supporting this idea, recent work shows that mast cells produce IL13 and drive PM maturation in autoimmune gastritis (AIG) ^14^. Whether ILC2s and mast cells play distinct or overlapping roles in *H. pylori* infection and AIG remains an open question.

We demonstrate that ILC2s play a key role in driving sex differences in the gastric immune response to *H. felis* infection. However, we also show that gastric T cells express the *Ar.* Androgens are well known to regulate T cell function, impairing effector responses against pathogens and promoting T cell exhaustion in tumors ^52-55^. Additionally, we identify sex differences in CD4/CD8 T cell ratios, with females exhibiting increased CD8+ T cell infiltration, which may indicate enhanced anti-tumor immune surveillance and contribute to the lower incidence of gastric cancer in females. While this study does not directly examine the role androgens in modulating T cell responses to *H. pylori,* our findings highlight the importance of future studies, as T cells are key effectors of *H. pylori-*driven gastric atrophy ^26^. In addition, our study does not address the effects of estrogens, which have been shown to enhance T cell activity and cytokine production, potentially increasing gastric epithelial damage during *H. pylori* infection ^56^. Notably, estrogens have also been reported to protect against *H. pylori*-driven gastric cancer in the hypergastrinemic INS-GAS model ^57, 58^. Further research is needed to fully elucidate the impact of steroid hormones on sex differences in *H. pylori* pathogenesis.

In summary, this study underscores the importance of investigating sex differences in *H. pylori* pathogenesis and gastric cancer. Our findings reveal a novel role for androgens in regulating gastric inflammation through an ILC2–APC–T cell axis. We demonstrate that androgen depletion exacerbates *Helicobacter* pathogenesis and accelerates preneoplastic lesion development. These results suggest that circulating androgen levels may serve as a useful prognostic biomarker for identifying men at increased risk of *H. pylori*-associated gastric cancer.

## Abbreviations

DHT: Dihydrotestosterone
ILC2: type two innate lymphoid cell
PM: pyloric metaplasia
APC: antigen presenting cell
DEG: differentially expressed gene
TCR: T cell receptor
TCRβδ KO: T cell receptor beta/delta knock out
ILC2-DTx: type two innate lymphoid cell diphtheria toxin depletion.

## Author Contributions

B.C.D. and J.T.B planned the study. B.C.D., M.T.M., J.L.P., S.K., and J.T.B performed experiments and analyzed data. B.C.D., L.W. and G.H. analyzed genomics data. B.C.D. and J.T.B analyzed all data and drafted the manuscript. All authors reviewed and approved the final manuscript.

## Acknowledgments

The authors acknowledge Biorender for facilitating the creation of the graphical abstract used in this report.

