## Supplementary Figures for "Androgen Signaling in Type 2 Innate Lymphoid Cells Drives Sex Differences in *Helicobacter*-Induced Gastric Inflammation and Atrophy"

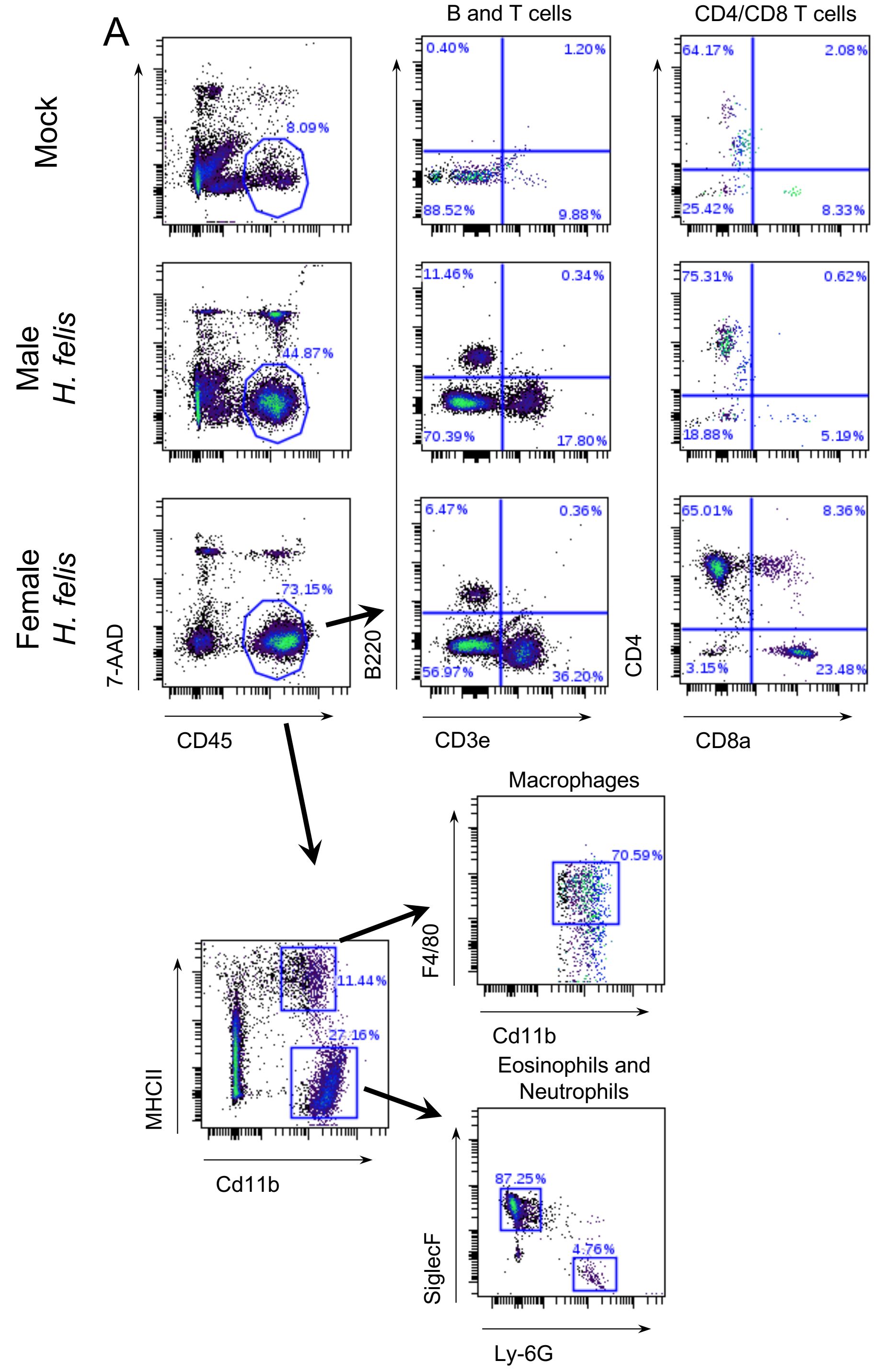


**Supplementary Figure 1:** Flow cytometry analysis of immune cell populations in male and female mice infected with *Helicobacter felis*. Panel A depicts the gating strategy and subsequent analysis for CD45+ leukocytes in mock (uninfected) and *H. felis*-infected male and female mice. The leftmost plots show the total CD45+ cell population. Middle plots demonstrate the segregation of B and T cells, and the CD4+/CD8+ T cell subsets. The rightmost plots provide a detailed analysis of macrophage identification through CD11b and F4/80 expression, and further classification of eosinophils and neutrophils using Siglec-F and Ly-6G markers, respectively.

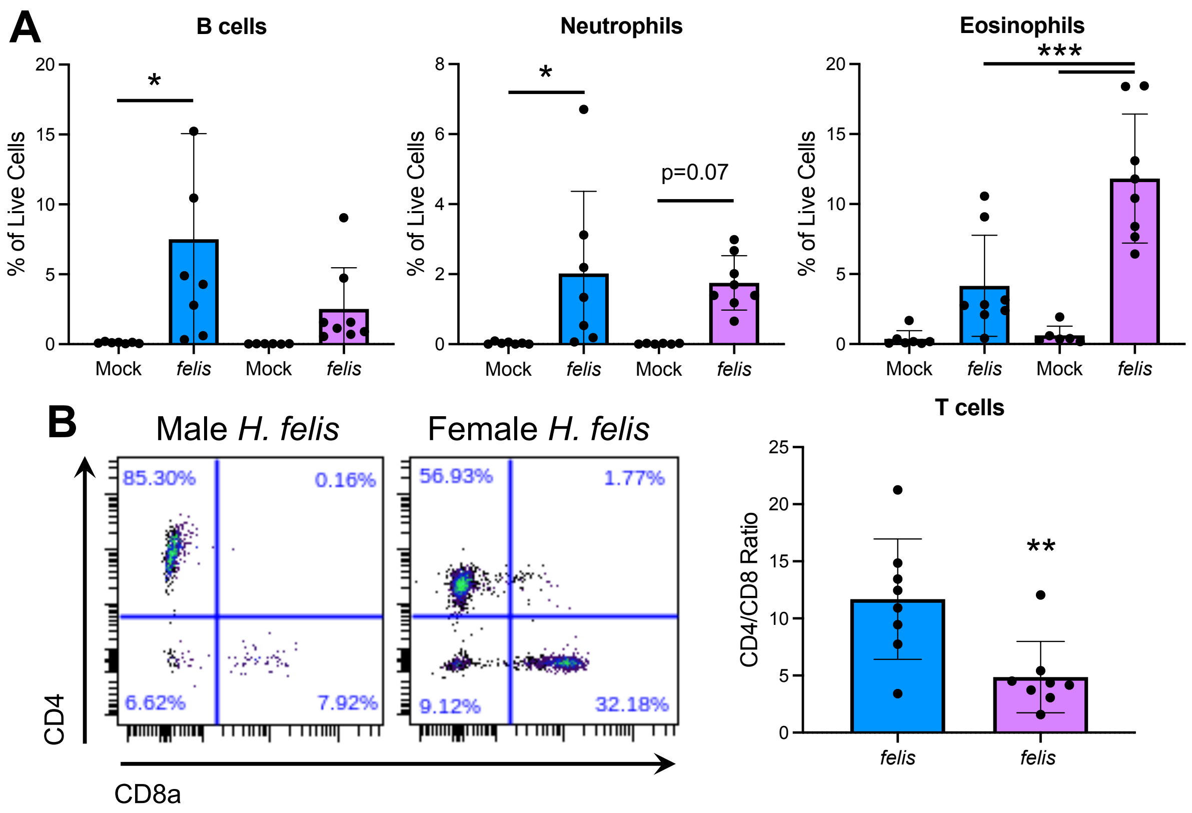


**Supplementary Figure 2:** Quantitative and phenotypic analysis of immune cell subsets in mock and *Helicobacter felis* infected male and female mice. (A) Quantification of B cells, neutrophils, eosinophils. (B) Representative flow cytometry plots showing CD4 and CD8a staining in T cells from male and female mice infected with *H. felis*. Plots highlight the differences in CD4/CD8 T cell ratios between the sexes. (*p < 0.05, **p < 0.01, ***p < 0.001). n≥6.

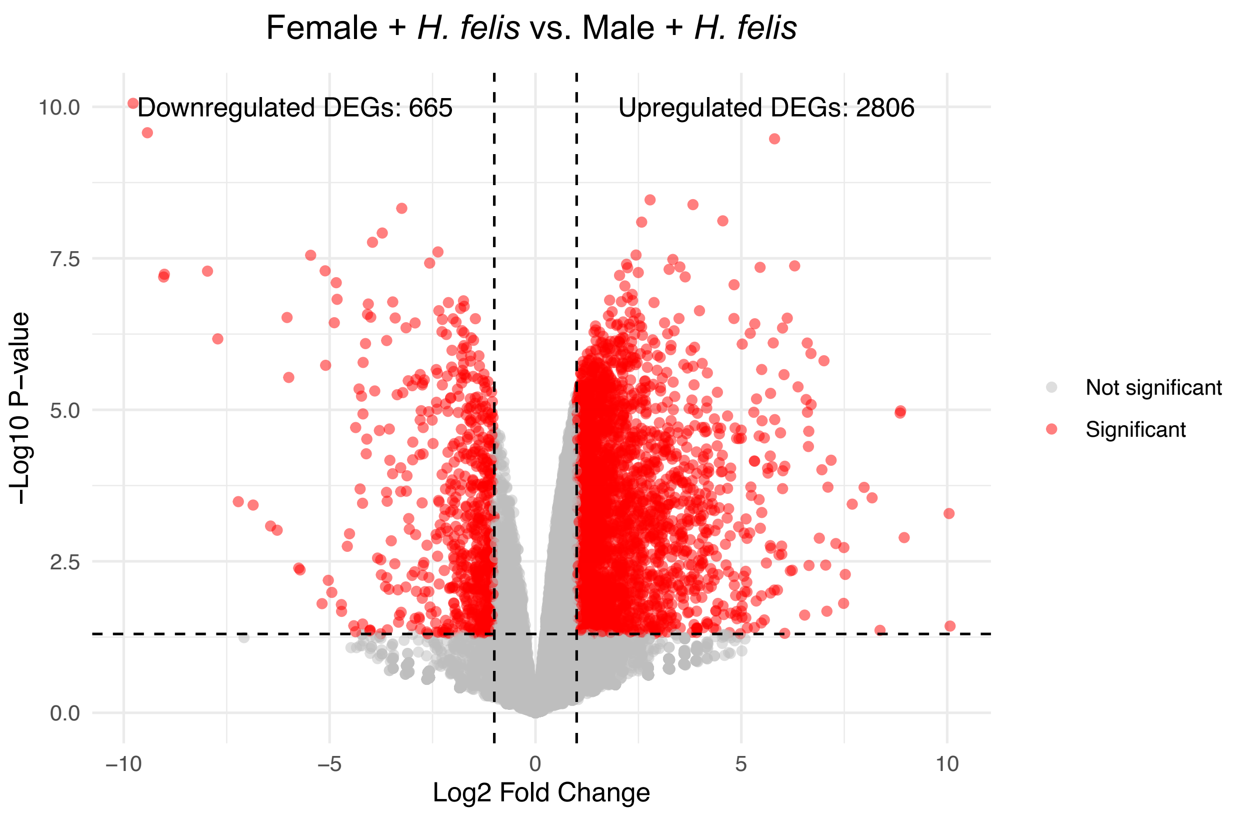


**Supplementary Figure 3:** Volcano plot illustrating differential gene expression in female versus male mice infected with *Helicobacter felis*. This plot compares the expression profiles between female and male mice, highlighting genes significantly upregulated (right) and downregulated (left) in females. Horizontal dashed line denotes the threshold for significance (-log10 p-value), and vertical dashed lines delineate the fold change boundaries. n=3.

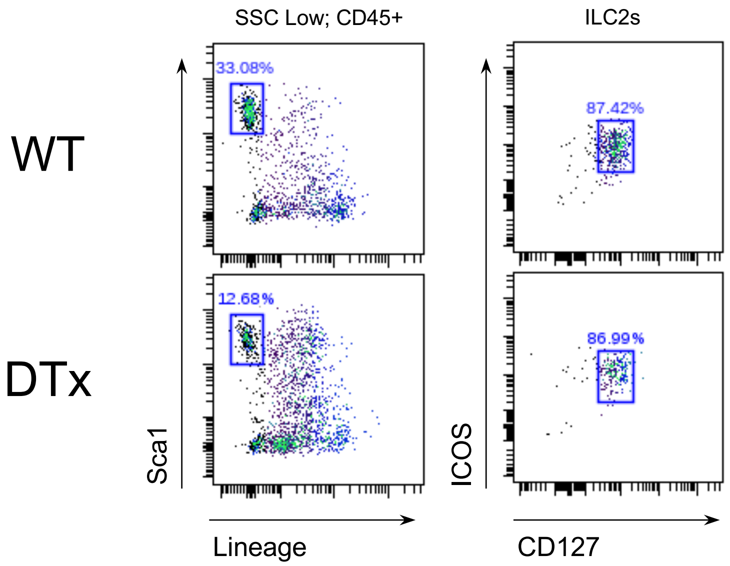


**Supplementary Figure 4.** Gating strategy used to identify gastric ILC2s. The top is a representative scatter plot of ILC2s in an uninfected WT mouse while bottom depicts a representative ILC2-DTx mouse.

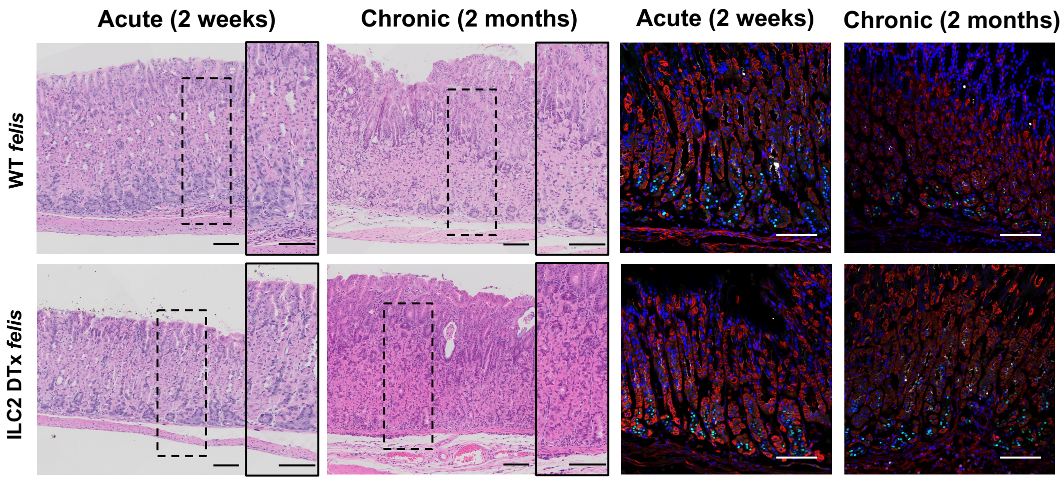


**Supplementary Figure 5.** Representative H&E and immunostaining micrographs of the gastric corpus from wild type (WT) and ILC2-DTx deficient male mice infected with *H. felis* for 2 weeks or 2 months. Immunofluorescent staining of corpus tissue probed with H+/K+ ATPase (red, parietal cells), MIST1 (green, chief cells), and GSII lectin (white, mucous neck cells) in the gastric mucosa. Nuclei are counterstained with DAPI (blue). Scale bars: 100 µm. n≥5.

Table 1: Metaplasia Transcripts

| Metaplasia Transcript | Publication PMID # |
| --- | --- |
| *Aqp5* | 34455107 |
| *Cd44* | 23848514 |
| *Cftr, Clu* | 22773549 |
| *Dmbt1* | 22944598 |
| *Gif* | 15647607 |
| *Gkn3* | 32835664 |
| *Muc6* | 10070955 |
| *Muc4* | 27125972 |
| *Olfm4* | 20398667 |
| *Tff2* | 10378506 |
| *Wfdc2* | 18242217 |
| *Pigr* | 24694107 |
| *Sox9* | 37270061 |
| *Chil4* | 31481545 |

**Supplementary Table 1.** Metaplastic transcripts identified from literature.
